# Hybrid correction of highly noisy Oxford Nanopore long reads using a variable-order de Bruijn graph

**DOI:** 10.1101/238808

**Authors:** Pierre Morisse, Thierry Lecroq, Arnaud Lefebvre

## Abstract

**Motivation:** The recent rise of long read sequencing technologies such as Pacific Biosciences and Oxford Nanopore allows to solve assembly problems for larger and more complex genomes than what allowed short reads technologies. However, these long reads are very noisy, reaching an error rate of around 10 to 15% for Pacific Biosciences, and up to 30% for Oxford Nanopore. The error correction problem has been tackled by either self-correcting the long reads, or using complementary short reads in a hybrid approach, but most methods only focus on Pacific Biosciences data, and do not apply to Oxford Nanopore reads. Moreover, even though recent chemistries from Oxford Nanopore promise to lower the error rate below 15%, it is still higher in practice, and correcting such noisy long reads remains an issue.

**Results:** We present HG-CoLoR, a hybrid error correction method that focuses on a seed-and-extend approach based on the alignment of the short reads to the long reads, followed by the traversal of a variable-order de Bruijn graph, built from the short reads. Our experiments show that HG-CoLoR manages to efficiently correct Oxford Nanopore long reads that display an error rate as high as 44%. When compared to other state-of-the-art long read error correction methods able to deal with Oxford Nanopore data, our experiments also show that HG-CoLoR provides the best trade-off between runtime and quality of the results, and is the only method able to efficiently scale to eukaryotic genomes.

**Availability and implementation:** HG-CoLoR is implemented is C++, supported on Linux platforms and freely available at https://github.com/morispi/HG-CoLoR

**Supplementary information:** Supplementary data are available at *Bioinformatics* online.

## 1 Introduction

Since a few years, long read sequencing technologies are being developed, and allow the solving of assembly problems for large and complex genomes that were, until then, hard to solve with the use of short reads sequencing technologies alone. The two major actors of these long read sequencing technologies are Pacific Biosciences and Oxford Nanopore. The latter, with the release of the MinION device, that can be run from a simple laptop, allows a low-cost and easy long read sequencing.

Even though long reads can reach lengths of tens of kbps, they also reach a very high error rate of around 10 to 15% for Pacific Biosciences, and up to 30% for Oxford Nanopore. Due to this high error rate, correcting long reads before using them in assembly problems is mandatory. Many methods are available for short read error correction, but these methods are not applicable to long reads, on the one hand because of their much higher error rate, and on the other hand, because most of the error correction tools for short reads focus on substitution errors, the main error type in Illumina data, whereas insertions and deletions are more frequent in long reads.

### 1.1 Related works

Recently, several methods for long read error correction have been developed. These methods can be divided into two main categories: either the long reads are self-corrected by aligning them against each other (PBDAG-Con (Chin *et al*., 2013), PBcR (Berlin *et al*., 2015)), or either a hybrid strategy, using complementary short reads is adopted. In this case, the short reads can either be aligned to the long reads (Nanocorr (Goodwin *et al*., 2015), CoLoRMap (Haghshenas *et al*., 2016)), or be assembled into contig on which the long reads are aligned (HALC (Bao and Lan, 2017)). de Bruijn graph based methods, where the long reads are corrected by traversing the paths of the graph, also started to develop recently, in the hybrid case (LoRDEC (Salmela and Rivals, 2014), Jabba (Miclotte *et al*., 2016)), as well as in the non-hybrid case (LoRMA (Salmela *et al*., 2017), Daccord (Tischler and Myers, 2017, unpublished)). NaS (Madoui *et al*., 2015), instead of using short reads to correct the long reads, uses the long reads as templates in order to recruit short reads and assemble them into contigs, used as corrected sequences. This approach requires to align the short reads to the long reads, in order to find seeds. The seeds are then compared to all the other short reads, in order to recruit new short reads, corresponding to low quality regions of the long read.

### 1.2 Limitations of current methods

Most of the current long read error correction methods only focus on Pacific Biosciences long reads. Therefore, they manage to perform error correction on long reads that display a maximum error rate of about 10 to 15%, but often do not manage to reduce the error rate at all when correcting long reads having higher error rates. As Oxford Nanopore, even with recent chemistries, faces difficulties lowering the error rate of the long reads below 15%, only a handful of methods can be applied on such data.

Moreover, those few methods, although managing to perform satisfying error correction, tend to yield unsatisfying assembly results. Our experiments show that only NaS manages to correct the long reads well enough so that they can assemble into a decent number of contigs, even on highly noisy data, but suffers from large runtimes.

### 1.3 Contribution

We introduce HG-CoLoR, a new long read hybrid error correction method that combines both the main idea from NaS to initiate the correction by using short reads that align to the long reads as seeds, and the use of a variable-order de Bruijn graph, built from the short reads, in order to get rid of the time consuming step of comparing all the short reads against each other. HG-CoLoR indeed focuses on an approach where the seeds are used as anchors on the variable-order de Bruijn graph, that is traversed in order to link them together and to produce the corrected long reads. Our experiments show that, while producing comparable results, even on highly noisy long reads, HG-CoLoR is several orders of magnitude faster than NaS. They also show that, when compared to state-of-the-art hybrid and non-hybrid long read error correction methods, HG-CoLoR provides the best trade-off between runtime and quality of the results, both in terms of reduction of the error rate and in terms of contiguity of the assemblies generated from the corrected long reads.

## 2 NaS Overview

NaS is a hybrid method for the error correction of long reads that, unlike other methods, generates corrected sequences from assemblies of short reads, instead of using the short reads to correct the long reads. More precisely, a corrected long read is produced as follows.

First, the short reads are aligned to the long read using BLAT (Kent, 2002) in fast mode, or LAST (Kielbasa *et al*., 2011) in sensitive mode, in order to find seeds, which are short reads that align correctly to the long read. Then, the discovered seeds are compared to all the other short reads with the help of Commet (Maillet *et al*., 2014), and similar short reads, which share a certain number of non-overlapping *k*-mers with the seeds, are recruited. Finally, the obtained subset of short reads is assembled using Newbler (from Roche company, unpublished), and a contig is produced, and used as the correction of the original long read.

Usually, a single contig is produced, but in repeated regions, a few bad reads can be recruited and yield erroneous contigs that must not be associated with the long read. To address this issue, and produce a single contig from multiple ones, NaS explicitly builds the contig-graph, and weights each node with the seeds coverage of the associated contig. Once the graph is built, the path with the highest total weight is chosen with the Floyd-Warshall algorithm, and contigs along that path are assembled to generate the final, unique contig. Finally, the short reads are aligned to the produced contig in order to verify its consistency. The contig is output and used as the correction of the initial long read if it is sufficiently covered by the short reads.

The reads recruitment is the most crucial step of the method, as it allows to retrieve short reads corresponding to low quality regions of the long read. However, this step is also the bottleneck of the whole NaS pipeline, as it is responsible for 70% of the total runtime on average.

## 3 Variable-order de Bruijn graph

### 3.1 de Bruijn graphs

The de Bruijn graph is a data structure that is widely used in assembly tools. Its nodes are defined as the fc-mers of the reads, and its edges represent prefix-suffix overlaps of length *k* – 1 between the fc-mers represented by the nodes. However, despite its usefulness, it is known that the Bruijn graph faces difficulties, due to the fact it fixes the *k*-mer size at construction time. On the one hand, choosing a high value of *k* will allow the graph to better deal with repeated regions, but will lead to missing edges in regions with locally insufficient coverage. On the other hand, choosing a small value of *k* will allow to correctly retrieve the edges of the graph in insufficiently covered regions, but will lead to more difficulties with repeated regions.

To overcome these problems, modern assemblers usually build multiple de Bruijn graphs of different orders. Although this approach allows to increase the quality of the produced assemblies, it also greatly increases both runtime and memory consumption, as multiple graphs need to be built, instead of a single one.

More recently, a few methods were developed to allow the representation of all the de Bruijn graphs, up to a maximum order *K*, in a single data structure. The manifold de Bruijn graph (Lin and Pevzner, 2014), for example, associates arbitrary substrings with nodes, instead of associating *k*-mers. This structure is however mainly of theoretical interest, as it has not been implemented yet. Another implementation of a variable-order de Bruijn graph has been proposed by Boucher *et al*. (2015). It relies on the succinct representation of the de Bruijn graph by Bowe *et al*. (2012), and supports additional operations that allow to change the order of the graph on the fly. However, the current implementation only supports construction up to an order of 64, which is too restrictive, as we do not want to limit the highest possible value of *K*.

To overcome this issues, we introduce a new implementation of the variable-order de Bruijn graph. It relies on PgSA (Kowalski *et al*., 2015), an index structure that allows to answer various queries on a set of reads.

### 3.2 PgSA overview

PgSA is a data structure that allows the indexing of a set of reads, in order to answer the following queries, for a given string *f*:

1. In which reads does *f* occur?
2. In how many reads does *f* occur?
3. What are the occurrences positions of *f*?
4. What is the number of occurrences of *f*?
5. In which reads does *f* occur only once?
6. In how many reads does *f* occur only once?
7. What are the occurrences positions of *f* in the reads where it occurs only once?

In these queries, *f* can be given either as a sequence of DNA symbols, or as a pair of numbers, representing respectively a read ID, and the starting position of *f* is this read.

As previously mentioned, in order to answer these queries, an index of the reads has to be built. PgSA builds it as follows. First, all reads with overlaps are concatenated with respect to these overlaps, in order to obtain a pseudogenome. If some reads for which no overlaps have been found are left after the pseudogenome creation, they are simply concatenated at the end of it. Then, a sparse suffix array of the pseudogenome is computed, along with an auxiliary array allowing the retrieval of the reads from the original set in the pseudogenome. Each record of this auxiliary array associates a read ID in the original set of reads to a read offset in the pseudogenome, and also contains flag data that bring complementary information about the read and that is used in order to handle the queries. The queries are processed by a simple binary search over the suffix array, coupled with the use of this complementary information.

As the reads are overlapped during the pseudogenome computation, and as PgSA does not record any information about their lengths, it only allows the indexing and querying of a set of reads of constant length. However, the length of the query string is not set at compilation time, and PgSA therefore supports queries for strings *f* of variable length.

### 3.3 Variable-order de Bruijn graph representation

A maximum order *K* is chosen, and the *K*-mers of the reads are indexed with PgSA, to be able to represent the nodes of all the de Bruijn graphs up to this maximum order. The edges of a given node, for any de Bruijn graph of order *k* ≤ *K*, are retrieved by querying the index, using the third query (*i.e*. what are the occurrences positions of *f*?), with the suffix of length *k* – 1 of the *k*-mer represented by the node. The query returns a set of pairs of numbers, each of these pairs representing respectively a *K*-mer ID and the occurrence position of the query string in that *K*-mer. The pairs are then processed, and only those whose position component does not represent the suffix of length *k* – 1 of the associated *K*-mer are retained (so that the occurrence can be extended to the right into a *k*-mer). These extended occurrences define the *k*-mers that have a prefix-suffix overlap of length *k* – 1 with the *k*-mer represented by the currently considered node, and thus the edges of this node.

As the edges are retrieved by querying the index, it is also easy to traverse the graph backward. For a given order *k*, instead of being queried with suffixes of the *k*-mers represented by the nodes, the index is simply queried with their prefixes. The returned sets of pairs are then processed in the same fashion as for forward traversal, except that only the pairs whose position component does not represent the prefix of length *k* – 1 of the associated *K*-mer are retained to define the edges. For better understanding, the algorithm allowing to retrieve the edges of any given node, forward or backward, in the de Bruijn graph of any order *k* ≤ *K*, is given in Algorithm 1.

## 4 Methods

### 4.1 Overview

HG-CoLoR, like NaS, aims to initiate the correction by using short reads that align to the long reads as seeds. However, its main objective is to get rid of the time consuming step of reads recruiting, that requires to compare the seeds to all the other short reads. To do so, it focuses on a seed-and-extend approach, and extended with the help of the previously described variable-order de Bruijn graph. This graph is built from the short reads, by choosing a maximum order *K* and indexing their *K*-mers with PgSA, and is traversed by querying the index, as previously described. For each long read, the graph is traversed in order to link together the associated seeds, **Algorithm 1**: Retrieve the edges of a given node. getOccurrencesPositions and getKmer are both PgSA functions that allow respectively to retrieve the occurrences positions of the given string in the set of *K*-mers, and to retrieve the sequence corresponding to the *K*-mer of identifier id. Line 2: Start with an empty set of edges. Lines 3–7: If traversing the graph forward, get the occurrences positions of the suffix of s in the set of *K*-mers, if traversing it backward, get the occurrences positions of its prefix. Lines 8–15: Process the list of occurrences positions. The processing is stopped when all the occurrences have been processed or when 4 edges have been found, as we work with the DNA alphabet and cannot find more than 4 edges per node. Lines 11–12: If traversing forward and if the position component does not represent the suffix of length *k* – 1 of the *K*-mer of identifier id, add an edge to the *k*-mer starting at position pos in this *K*-mer. Line 13–14: If traversing backward and if the position component does not represent the prefix of length *k* – 1 of the *K*-mer of identifier id, add an edge to the *k*-mer starting at position pos – 1 in this *K*-mer. used as anchors. The path of the graph that was followed to link two seeds together thus dictates a corrected sequence for the missing part of the long read. Finally, once all the seeds have been linked, the tips of the obtained sequence are extended by traversing the graph again, to reach the borders of the original long read. HG-CoLoR’s workflow is summarized in Figure 1, and its four main steps are described below.

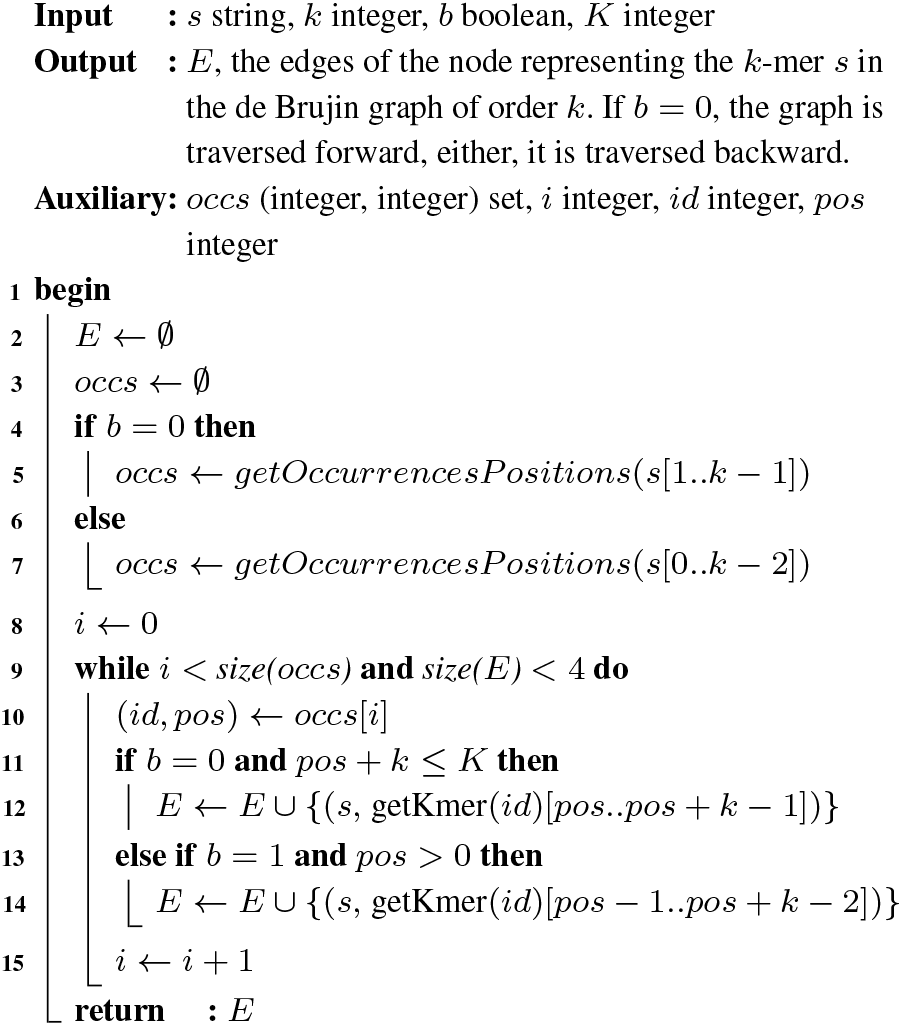

**Fig. 1.**
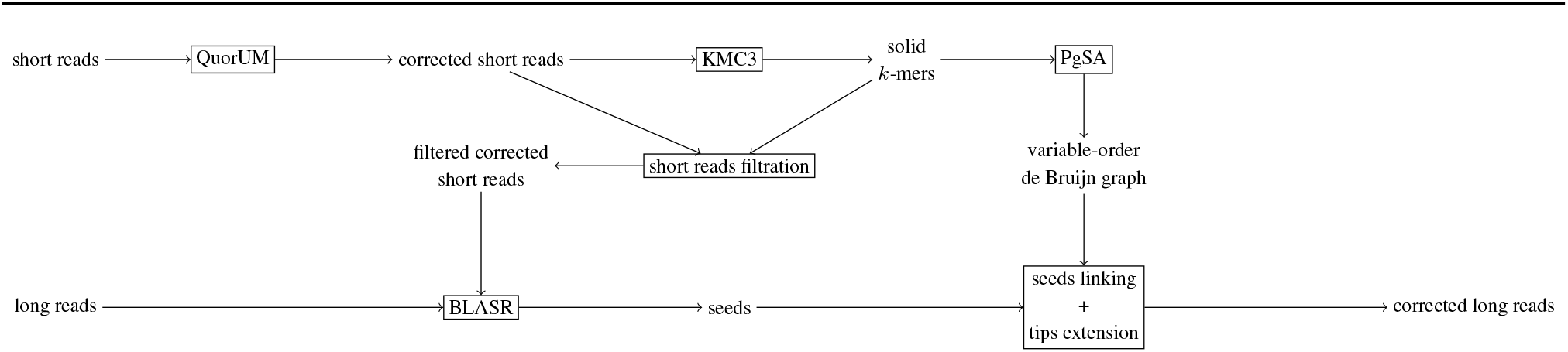
HG-CoLoR’s workflow. First, the short reads are corrected with QuorUM in order to get rid of as much sequencing errors as possible. Then, a maximum order *K* is chosen for the graph, and the *K*-mers from the corrected short reads are obtained with KMC3. To further reduce the error rate, a filtration step is applied to the corrected short reads, and those containing weak *K*-mers areremoved. Forthe samereason, onlythe solid *K*-mers fromthe corrected shortreads are indexed with PgSA, to representthe variable-order de Bruijn graph. The previously filtered corrected short reads are then aligned to the long reads with the help of BLASR in order to find seeds. Each long read is then processed independently. For each of them, the graph is traversed in order to link together the associated seeds, used as anchors, in order to retrieve corrected sequences for the missing parts of the long read. Then, the tips of the sequence obtained after linking together all the seeds are extended in both directions by traversing the graph, to reach the initial long read’s borders. Finally, the corrected long read is output.

Despite high similarities with other graph based methods, in particular with LoRDEC, using short reads that align to the long reads as anchors on the graph, is quite different from using solid *k*-mers from the long reads. Indeed, in the case of Oxford Nanopore data, due to the very high error rate of the long reads, even short, solid *k*-mers have a high chance to be erroneous. Such erroneous *k*-mers would therefore lead to the use of bad anchors, and thus to unsatisfying correction results. However, as short reads are accurate, they can be used as reliable anchors, with little to no chance of being erroneous. Moreover, using short reads as anchors also allows to directly build the graph with large values of *k*, without needing to perform multiple rounds of correction, increasing the value of *k* at each step, in the same fashion as LoRMA.

### 4.2 Short reads correction and graph construction

Even though short reads are already accurate prior to any correction, they still contain a small fraction of errors. As HG-CoLoR seeks to build a variable-order de Bruijn graph of high maximum order from the short reads, as much errors as possible have to be removed from this data, to avoid erroneous paths in the graph. To do so, the short reads are corrected with the help of QuorUM (Marçais *et al*., 2015), which provides the best trade-off between runtime and quality of the correction, among all the tested short read error correction tools.

A maximum order *K* is then chosen for the graph, and the *K*-mers from the corrected short reads are extracted with KMC3 (Kokot *et al*., 2017). To further reduce the error rate of the short reads data, and thus avoid bad seeds and chimeric paths on the graph, short reads containing weak *K*-mers (*i.e. K*-mers that appear less than a certain threshold) are filtered out and not used in the following steps, and only the solid *K*-mers are used to build the graph.

### 4.3 Seeds retrieving and merging

Like with NaS, the seeds are found by aligning the short reads to the long reads. This step is performed with thehelpofBLASR (ChaissonandTesler, 2012), an alignment tool originally designed to align long reads dominated by insertion and deletion errors to a reference genome, that however also manages to nicely deal with this type of errors when aligning short reads to long reads. Each long read is then processed independently, and two phases of analysis are applied to the associated seeds.

First, if the alignment positions of a given pair of seeds indicate that they overlap over a sufficient length, their assumed overlapping sequences are compared, and the two seeds are merged accordingly. If the alignment positions indicate that the two seeds do overlap, but over an insufficient length, or if the assumed overlapping sequences do not coincide, only the seed with the best alignment score is kept.

Second, once all the seeds with overlapping alignment positions have been merged or filtered out, sequence overlaps between consecutive seeds having close alignment positions are computed. As in the previous step, if a given seed perfectly overlaps another one over a sufficient length, the two seeds are merged. This step allows to take into account small insertion errors in the long reads that were not detected during the alignment step, and that could lead to difficult linkings in the next step.

### 4.4 Seeds linking

Once the seeds have been found and merged for all the long reads, HG-CoLoR processes each of the long reads independently and attempts to link together their associated seeds by considering them as pairs, and traversing the graph. For a given pair, the seed that has the leftmost alignment position is called the *source*, and the one that has the rightmost alignment position is called the *target*. To link a pair of seeds together, the rightmost *K*-mer of the source and the leftmost *K*-mer of the target are used as anchors on the graph. The graph is then traversed, in order to find a path between the two anchors. When such a path is found, the sequence it dictates is used as a correction for the missing part of the long read.

HG-CoLoR traverses the variable-order de Bruijn graph starting from its highest order. The order is decreased at a given node only if this node does not have any edge for the current order, or if all its edges for the current order have already been explored and did not allow to reach the destination. When the order of the graph is decreased, the size of the *k*-mers from the source and from the destination is decreased accordingly, so that they can still be used as anchors. A minimum order is also set, so that HG-CoLoR does not traverse de Bruijn graphs representing short, and probably meaningless overlaps. When facing branching paths for a given order *k*, HG-CoLoR performs a greedy selection. The edge leading to the node representing the *k*-mer having the highest number of occurrences is therefore explored first. This greedy selection allows to avoid traversing too many nodes representing *k*-mers having low frequencies, that, despite the correction and filtration steps, may contain a sequencing error. When a path from the source to the target is found, it is considered as optimal due to the greedy selection and to the fact that the order of the graph is only locally decreased. It is thus chosen as the correction for the missing part of the long read. We voluntarily select the optimal path this way, instead of exploring multiple ones and selecting the one that aligns the best to the long read as the correction, in order to avoid prohibitive runtimes.

We also set a mismatches threshold *t* when linking two seeds together. We therefore consider that the source and the target can be linked together if a path starting from the anchor *K*-mer of the source reaches a *K*-mer having less than *t* mismatches with the anchor *K*-mer of the target. Such a threshold allows to overcome the few mismatches errors that can still be present on the seeds, despite the correction and filtration steps.

However, short reads from a different region of the reference genome may align to the long read and may be used as seeds. As such seeds could lead to impossible linkings, even if all the existing paths of the graph were explored, a threshold on the maximum number of branches explorations is set. If this threshold is reached, and no path has been found to link the source and the target together, the current linking iteration is given up, and HG-CoLoR attempts to skip the target that could not be reached. In other words, the source remains the same, the target that could not be reached is ignored, the target is defined as the following seed, and a new linking iteration is performed. An illustration of this process is given in Supplementary Figure S2.

As skipping seeds can lead to an important number of failed linking attempts, if erroneous seeds are present in great proportion on a long read, a threshold on the maximum number of seeds that can be skipped is set. Once this threshold is reached, as none of the linking attempts succeeded, HG-CoLoR splits the corrected long read. The part corresponding to the seeds linked so far is output, and the graph is traversed again, in order to try to link together the remaining seeds, including the ones it attempted to skip, independently of the previous part. We chose to always split the long reads in order to avoid reporting erroneous bases from the long reads, or bases corresponding to wrongly aligned seeds. Reporting such bases would indeed decrease the quality of the correction, and negatively impact the downstream assemblies.

### 4.5 Tips extension

Finally, it is obvious that the seeds do not always align right at the beginning and at the end of the long reads. Thus, in order to get as close as possible to its original length, once all the seeds of a given long read have been linked, HG-CoLoR keeps on traversing the graph to extend the tips of the produced corrected long read. In the same fashion as in the previous step, the traversal starts from the highest order of the variable-order de Bruijn graph, and the order is decreased at a given node only if this node does not have any edge for the current order. The tips of the corrected long read are thus extended until either the original long read’s borders or a branching path are reached. Indeed, in the case of tips extension, when facing a branching path, HG-CoLoR has no clue as to which path to chose and continue the extension with, nor any anchors, unlike when it attempts to link two seeds together. Therefore, greedy selection and exploration of multiple branches are useless and the extension is simply stopped when such a situation occurs. In the case of split long reads, every fragment is extended as mentioned.

## 5 Results and discussion

We ran experiments on three real datasets of inscreasing size: one from *A. baylyi*, one from *E. coli* and one from *S. cerevisae*. They include respectively 381 Mbp, 134 Mbp and 1,173 Mbp of Oxford Nanopore MinION long reads, and224Mbp, 232Mbpand625 Mbpoflllumina short reads. All details are given in Supplementary Table S1. Unless otherwhise sepecified, all experiments were run on a 32 GB RAM machine equipped with 16 cores.

We compare HG-CoLoR against hybrid error correction tools NaS and Jabba, and also against two self-correction methods, namely Daccord and the method used in the assembler Canu (Koren *et al*., 2017). We evaluate the accuracy of the different tools with two different approaches. First, we analyze how well the long reads were corrected, by aligning them to the reference genomes, and second, we investigate the quality of the assemblies that can be generated from the corrected long reads. The Nanocorr, CoLoRMAP, LoRDEC, LoRMA, and HALC softwares were also tested, but as they led to unsatisfying results, we discard them from the comparison. Due to its large runtimes, NaS was only executed in fast mode.

### 5.1 Parameters

We ran multiple rounds of correction with HG-CoLoR on the *S. cerevisae* dataset to experiment with the parameters. Thereby, we found that using a variable-order de Bruijn graph of maximum order *K* = 100 yielded the best compromise between runtime, number of corrected long reads, proportion of split long reads, average length and number of corrected bases (see Supplementary Figure S1). The minimum overlap length to allow the merging of two seeds during the second step was set to 99, accordingly to the maximum order *K* chosen for the graph. The minimum order of the graph was set to *k* = 50, as setting it to larger values resulted in less corrected long reads, that were more split, and thus shorter, due to local drops of coverage. Setting it to smaller values also resulted in more split, and shorter long reads, due to the exploration of meaningless edges, especially in repeated regions, in addition to larger runtimes (see Supplementary Figure S2). The maximum number of branches explorations was set to 1,500, as decreasing it also resulted in more split, and shorter long reads, and increasing it more barely yielded better results, but increased the runtime (see Supplementary Figure S3). For similar reasons, the maximum number of seed skips was set to 5, and the mismatches threshold was set to 3. For the alignment of the short reads to the long reads, BLASR was used with default parameters except for bestn, that was set to 50 instead of 10, in order to obtain a greater number of seeds, and therefore correct more long reads. Yet again, increasing this parameter to larger values only impacted the runtime, and did not meaningfully improve the correction results, while decreasing it induced a drop of the number of corrected long reads. As we only use a 50x coverage of short reads, the *K*-mer solidity threshold was set to 1 (*i.e*. all the *K*-mers were considered as solid). Canu was run with parameters `-correct, -nanopore-raw, stopOnReadQuality=false`, due to the high error rate of the long reads, `corOutCoverage=300`, in order to correct as many long reads as possible, and genomeSize set to the exact number of bases of each reference genome. Other tools were run with default or recommended parameters. To allow better comparison, the short reads were corrected with QuorUM before running Jabba, instead of using Karect (Allam *et al*., 2015), the tool recommended by the authors. All tools were run with 16 processes.

### 5.2 Comparison of the quality of error correction

The long reads were aligned with Last prior to any correction, as it deals better with raw long reads. The different correction tools were then run, and the obtained corrected long reads were aligned with BWA mem (Li and Durbin, 2010) given their high accuracy. Results are given in Table 1 and discussed below.

**Table 1.** Statistics of the long reads, before and after correction by the different methods. The number of reads column account for the number of corrected long reads, not for the number of output fragments. Precise runtime is omitted for NaS on S. cerevisae because the results did not compute in 16 days, and the execution was therefore stopped. NaS corrected reads for this dataset were obtained from the Genoscope website. Results are omitted for the two selfcorrection tools on S.cerevisae, since they could not correct the long reads.

| Method | Original | Jabba | NaS | HG-CoLoR | Canu | Daccord |
| --- | --- | --- | --- | --- | --- | --- |
| *A. baylyi* |  |  |  |  |  |  |
| Number of reads | 89,011 | 16,618 | 24,063 | 25,214 | 8,122 | 19,623 |
| Split reads (%) | 0 | 4.90 | 0 | 0.82 | 5.47 | 53.02 |
| Average length | 4,284 | 10,260 | 8,840 | 11,400 | 9,345 | 3,244 |
| Number of bases (Mbp) | 381 | 179 | 213 | 290 | 81 | 175 |
| Average identity (%) | 70.09 | 99.40 | 99.82 | 99.73 | 97.79 | 91.92 |
| Genome coverage (%) | 100 | 99.82 | 100 | 100 | 99.79 | 100 |
| Runtime | N/A | 2min | 94h18min | 1h38min | 32min | 45min |
| *E. coli* |  |  |  |  |  |  |
| Number of reads | 22,270 | 21,005 | 21,818 | 21,961 | 17,154 | 17,478 |
| Split reads (%) | 0 | 4.98 | 0 | 0.10 | 0.38 | 34.40 |
| Average length | 5,999 | 5,797 | 7,926 | 6,115 | 7,080 | 4,495 |
| Number of bases (Mbp) | 134 | 128 | 173 | 134 | 122 | 119 |
| Average identity (%) | 79.46 | 99.81 | 99.86 | 99.71 | 96.23 | 98.51 |
| Genome coverage (%) | 100 | 99.43 | 100 | 100 | 99.99 | 99.99 |
| Runtime | N/A | 3min | 72h02min | 1h11min | 36min | 30min |
| *S. cerevisiae* |  |  |  |  |  |  |
| Number of reads | 205,923 | 33,484 | 71,793 | 70,305 | — | — |
| Split reads (%) | 0 | 11.47 | 0 | 5.80 | — | — |
| Average length | 5,698 | 6,455 | 5,938 | 6,961 | — | — |
| Number of bases (Mbp) | 1,173 | 243 | 426 | 521 | — | — |
| Average identity (%) | 55.49 | 99.54 | 99.59 | 98.96 | — | — |
| Genome coverage (%) | 99.90 | 93.32 | 98.70 | 99.31 | — | — |
| Runtime | N/A | 12min | > 16 days | 10h53min | — | — |

Jabba clearly performed the best when it comes to runtime, outperforming all the other tools by several orders of magnitude. It also produced corrected long reads that aligned with a high identity. However, although highly accurate, these corrected long reads did not manage to completely cover any of the reference genomes. These unresolved regions likely come from the important proportion of split long reads that were produced, due to the fact that Jabba uses a de Bruin graph of fixed order, and therefore faces problems with local drops of coverage. Pre-processing the short reads using Karect, as recommended by the authors, did not show any significant improvement (see Supplementary Table 2).

Apart from Jabba, the two self-correction tools outperformed the two others hybrid-correction methods in terms of runtime. However, the error correction was not very efficient, as the produced corrected long reads still displayed a large proportion of errors, as high as 8% for those produced by Daccord on the *A. baylyi* dataset. The average length of the long reads corrected with Daccord was also smaller than the average length of the original long reads, due to the high proportion of split long reads. Even on a cluster with large resources none of these two tools managed to correct the highly noisy *S. cerevisae* dataset, underlining the fact that hybrid error correction remains the only way to to correct highly noisy Oxford Nanopore long reads.

Therefore, only NaS and HG-CoLoR managed to produce corrected long reads that covered the whole reference genomes with a high identity, except for a few regions of *S. cerevisae*, due to the fact that neither the original long reads nor the short reads did cover the whole genome. Despite a smaller number of output corrected long reads, HG-CoLoR yielded more corrected bases than NaS, and covered the reference genome better. For all the datasets, the long reads corrected with NaS aligned with a slightly higher identity than those corrected with HG-CoLoR. However, despite being run in fast mode, NaS was several orders of magnitude slower than HG-CoLoR on all the datasets.

As a result, despite its larger runtimes than self-correction methods, and its slight disadvantage on the alignment identity of the corrected long reads when compared to NaS, HG-CoLoR displayed the best trade off between runtime and quality of the results.

### 5.3 Comparison of the quality of assembly

The corrected long reads were assembled using Canu, without the correction and trimming steps. The following parameters were used for all of the assemblies: `OvlMerSize=17, OvlMerDistinct = 0.9925, OvlMerTotal = 0.9925`. The `genomeSize` parameter was set independently to the exact number of bases of each reference genome. All the other parameters were set to their default values. Comparisons of the assemblies against the reference genomes were performed with MUMmer (Kurtz *et al*., 2004). Results are given in Table 2 and discussed below.

**Table 2.** Statistics of the assemblies generated from the corrected long reads. Reported identities stand for the 1-to-1 alignments.

| Method | Jabba | NaS | HG-CoLoR | Canu | Daccord |
| --- | --- | --- | --- | --- | --- |
| *A. baylyi* |  |  |  |  |  |
| Long reads coverage | 50x | 59x | 81x | 22x | 49x |
| Number of contigs | 14 | 1 | 1 | 3 | 1 |
| NG50 | 216,679 | 3,629,508 | 3,634,118 | 2,887,573 | 3,520,381 |
| Genome coverage (%) | 89.03 | 100 | 99.99 | 99.39 | 100 |
| Identity (%) | 99.94 | 99.99 | 99.94 | 97.04 | 97.06 |
| *E. coli* |  |  |  |  |  |
| Long reads coverage | 28x | 37x | 29x | 26x | 26x |
| Number of contigs | 41 | 1 | 1 | 3 | 2 |
| NG50 | 138,730 | 4,635,116 | 4,703,199 | 3,155,369 | 4,558,944 |
| Genome coverage (%) | 95.81 | 99.90 | 100 | 99.82 | 100 |
| Identity (%) | 99.99 | 99.99 | 99.99 | 97.23 | 97.84 |
| *S. cerevisiae* |  |  |  |  |  |
| Long reads coverage | 20x | 35x | 43x | — | — |
| Number of contigs | 138 | 119 | 77 | — | — |
| NG50 | 47,164 | 146,459 | 292,707 | — | — |
| Genome coverage (%) | 68.67 | 97.44 | 97.08 | — | — |
| Identity (%) | 99.99 | 99.95 | 99.89 | — | — |

In agreement with what we observed in Table 1, the fact that the long reads corrected with Jabba did not manage to cover the whole reference genomes resulted in highly fragmented assemblies, that could not resolve large regions of the reference genomes. As a result, despite their high accuracy, those corrected long reads yielded the most fragmented assemblies, that covered the least the reference genomes, and that displayed the smallest NG50 sizes.

Surprisingly, the long reads corrected with the two self-correction tools, despite their higher error rate and their weaker coverage depth than long reads corrected with the hybrid tools, assembled into a small number of contigs, that displayed high NG50 sizes, both on the *A. baylyi* and the *E. coli* datasets. However, due to the high error rate of the long reads, these assemblies displayed the lowest identities when compared to the reference genomes. Moreover, on these two datasets, assemblies generated from long reads corrected with Daccord outperformed those generated from long reads corrected with Canu in terms of contiguity, NG50 size and coverage of the reference genomes. On the *A. baylyi* dataset, the long reads corrected with Daccord even assembled into a single contig.

On the *A. baylyi* and the *E. coli* datasets, NaS and HG-CoLoR produced corrected long reads that assembled into a single contig. The NG50 sizes of these assemblies were highly similar, and the only differences between the two tools were a small region of *E. coli* that was not covered by the assembly generated from the long reads corrected with NaS, and a slightly lower identity when compared to the reference genome for the assembly generated from the long reads corrected with HG-CoLoR on the *A. baylyi* dataset. However, on the *S. cerevisae* dataset, the assembly generated from long reads corrected with HG-CoLoR outperformed the assembly generated from long reads corrected with NaS in terms of contiguity and NG50 size, even though it slightly less covered the reference genome.

### 5.4 Scalability

To investigate the scalability of our method, we tested it on a dataset from the larger eukaryotic genome of *C. elegans*. It includes 2 Gb of real Oxford Nanopore MinION long reads and 5 Gb of Illumina short reads simulated with ART (Huang *et al*., 2012), as no real Illumina reads of satisfying length quality were available. Details are given in Supplementary Table S1. NaS was not run due to its prohibitive runtime, and Daccord did not manage to perform correction, even on a cluster with large resources. We therefore only compare HG-CoLoR against Jabba and Canu. Error correction and assembly statistics are given in Table 3. Similarly to our previous observations, Jabba performed orders of magnitude faster than the two other tools, and produced high quality corrected long reads, that weakly covered the reference genome, and yielded an unsatisfying assembly. Canu also performed faster than HG-CoLoR, and produced more corrected long reads, that covered the reference genome well, but still displayed a high proportion of error. The assembly generated from these reads thus displayed a low identity, and, surprisingly, failed to resolve 11% of the reference genome. HG-CoLoR managed to correct the long reads so that they both display a high identity and cover the reference genome well. The assembly generated from these reads also had a high identity, and displayed the highest proportion of genome coverage. The runtime was similar to NaS on the *A. baylyi* dataset, that contained close to 12 times less bases, attesting the better scalability of our method. Moreover, it is worth noting that the memory peak for HG-CoLoR was only of 10GB, making it able to scale to large genomes even on a reasonable setup.

**Table 3.** Statistics of the long reads and of the generated assemblies on the C. elegans dataset. Omitted NG50 sizes mean that the assemblies did not reach half of the genome size.

| Method | Original | Jabba | HG-CoLoR | Canu |
| --- | --- | --- | --- | --- |
| **Correction** |  |  |  |  |
| Number of reads | 363,500 | 219,925 | 271,738 | 340,826 |
| Split reads (%) | 0 | 20,46 | 11.63 | 0 |
| Average length | 5,524 | 3,910 | 5,161 | 5,408 |
| Number of bases (Mbp) | 2,008 | 1,060 | 1,599 | 1,843 |
| Average identity (%) | 71.07 | 99.85 | 98.95 | 85.63 |
| Genome coverage (%) | 99.99 | 95.43 | 99.88 | 99.89 |
| Runtime | N/A | 1h16min | 94h21min | 16h38min |
| **Assembly** |  |  |  |  |
| Long reads coverage | 20x | 11x | 16x | 18x |
| Number of contigs | 530 | 1,374 | 830 | 1,049 |
| NG50 | — | — | 183,098 | 116,510 |
| Genome coverage (%) | 15.80 | 53.62 | 94.91 | 88.75 |
| Identity (%) | 92.99 | 99.97 | 99.87 | 95.60 |

## 6 Conclusion

We described HG-CoLoR, a new hybrid method for the error correction of long reads, that, like NaS, initiates the correction by using short reads that align to the long reads as seeds. HG-CoLoR however, instead of recruiting new short reads in an all-against-all comparison step and assembling them, like NaS, relies on an extension step based on a variable-order de Bruijn graph. This graph, which is built from the short reads, is used to extend and link together the seeds, which are used as anchors, in order to correct uncovered regions of the long reads by a simple traversal.

Our experiments show that, compared against state-of-the-art hybrid and non-hybrid error correction tools HG-CoLoR, offers the best trade-off between runtime and quality of the results, both in terms of quality of the error correction itself, and in terms of quality of the assemblies generated from the corrected long reads. Further experiments also show that our method is the only one able to efficiently scale to eukaryotic genomes.

The development of this method and our experiments underline the fact that, despite already being useful, self-correction methods are still not completely applicable to Oxford Nanopore long reads. Indeed, they do not manage to perform error correction at all on long reads sequenced with early chemistries, that display a very high error rate. They also do not scale to eukaryotic genomes, either completely failing to perform correction, or barely reducing the error rate, despite an acceptable error rate of the original long reads. Therefore, hybrid approaches remain interesting to correct Oxford Nanopore long reads, either in the case of large genomes, or in the case of very high error rates, as resequencing with more recent chemistries is not always affordable, and that even recent sequencings rarely display an error rate below 15% in practice.

As further work, we plan to focus on a new implementation of PgSA, as the current one does not support parallel querying of the index, and therefore forces current implementation of HG-CoLoR to use mutexes. Getting rid of that need would reduce the runtime of the method. Another index structure allowing to query a set of *k*-mers with strings of variable length, and supporting parallel querying, could also replace PgSA. Another direction is to try out other aligners for the alignment step of the short reads to the long reads, in order to possibly discover the seeds quicker, or correct more long reads.

## Acknowledgements

The authors would like to thank the Genoscope for the availability of most of the data used in this paper. Part of the computation for this work has been executed on intensive computation resources from the CRIANN.

